# Single-cell trajectories in metastatic urothelial carcinoma reveal tumor–immune reprogramming and macrophage-driven resistance to PD-(L)1 blockade

**DOI:** 10.64898/2026.03.31.715549

**Authors:** Ronan Flippot, Amélie Roehrig, Julien Vibert, Nicolas Stransky, Luc Cabel, Kevin Mulder, Benjamin Besse, Claudio Nicotra, Maud Ngo Camus, Christophe Massard, Etienne Rouleau, Gerome Jules-Clement, Alicia Tran-Dien, Lambros Tselikas, Constance Thibault, Mostefa Bennamoun, Gromoslaw A. Smolen, Mukund Varma, Ruth Kulicke, Jean-Yves Scoazec, Céline Vallot, Maud Kamal, Agathe Peltier, Eric Letouzé, Yohann Loriot

## Abstract

Immune checkpoint inhibitors (ICI) improved outcomes in metastatic urothelial carcinoma (mUC), but primary and acquired resistance remain poorly understood. We performed single-nuclei RNA sequencing on sequential metastatic biopsies from ICI-treated mUC patients. Tumor cells showed transcriptomic heterogeneity within individual lesions, basal cells being associated with increased immune infiltration and response. Myeloid and lymphoid compartments exhibited features of immune dysfunction in non-responders. Longitudinal analyses revealed convergent adaptive resistance mechanisms, dominated by polarization toward pro-tumoral macrophage states, but also including downregulation of the antigen presentation machinery in tumor cells, increased checkpoint expression with loss of cytotoxicity in T cells. Individual trajectories point to distinct evolutionary routes under ICI pressure. Across pivotal ICI trials, bulk expression of the M2-like macrophage marker HES1 predicted ICI resistance. Our study provides the first single-cell longitudinal atlas of ICI-treated mUC, revealing macrophage reprogramming as a dominant driver of resistance, establishing a framework for individualized immunotherapy strategies.

## Introduction

Metastatic urothelial carcinoma (mUC) remains a lethal disease despite recent therapeutic advances^1^. Immune checkpoint inhibitors (ICIs) targeting PD-L(1) have improved survival across treatment settings, yet only a minority of patients derive durable benefit^2–6^. Most individuals experience primary resistance, and among initial responders, acquired resistance is almost universal. While PD-L1 immunohistochemistry and common genomic alterations offer limited predictive value in metastatic setting, the biological determinants that shape sensitivity or resistance to ICIs in mUC remain largely undefined.

Several studies in localized muscle-invasive bladder cancer (MIBC) have suggested the importance of pre-existing immune contexture, including T and B cell infiltration, tertiary lymphoid structures and inflammatory transcriptional programs^7–13^. However, these analyses mostly relied on bulk transcriptomics or immunohistochemistry from primary tumors, which may not reflect the biology of metastatic lesions. Bulk-based analyses also obscure the cellular heterogeneity of both tumor and immune compartments, preventing refined characterization of antitumor versus protumoral states. Single-cell technologies have begun to reveal the complexity of urothelial carcinoma, including luminal-basal plasticity and diverse myeloid cell states, but these studies remains cross-sectional and do not capture the dynamic rewiring induced by ICIs.

Understanding how the tumor and its microenvironment evolve over time during immunotherapy is critical to unravel resistance mechanisms. In solid tumors, emerging evidence indicates that ICI resistance may arise through multiple non-mutually-exclusive processes, including antigen presentation defects (HLA or B2-microglobulin loss), impaired IFN signaling, T cell exhaustion and expansion of immunosuppressive myeloid populations. However, whether these mechanisms operate in metastatic urothelial carcinoma, how they evolve under selective pressure, and how they combine within individual patients remain unknown. No study to date has mapped, at the single-cell level the longitudinal trajectories of tumor, lymphoid and myeloid compartments in mUC during PD-L(1) blockade.

To address this gap, we conducted an in-depth, single-cell analysis of metastatic tumor biopsies in the MATCH-R biological trial (NCT02517892) which prospectively collected sequential metastatic samples at baseline, during therapy and at progression in patients treated with anti-PD-(L)1 monotherapy. Using single-nuclei RNA sequencings supported by whole-exome sequencing and bulk transcriptomics, we generated a dataset of more than 220,000 high-quality cells mapped across 55 biopsies. This design provides a unique opportunity to dissect both pre-treatment determinants of response and adaptive resistance mechanisms emerging under immune pressure.

Our study reveals that mUC lesions display extensive intratumoral transcriptomic heterogeneity, including co-occurrence of basal, luminal and neuroendocrine-like programs within individual samples. We identify tumor-intrinsic and microenvironmental features associated with ICI response, including the proportion of basal tumor cells and distinct CD4/CD8 T-cell states. Importantly, we uncover the central role of macrophage reprogramming with enrichment of HES1+ M2-like immunosuppressive states predicting both primary resistance and subsequent progression. Longitudinal analyses show convergent adaptive programs across patients such as downregulation of antigen-presentation machinery and IFN signaling in tumor cells, increased immune checkpoint expression and exhaustion in T cells and dominant polarization toward protumoral macrophage subsets. These mechanisms combine in patient-specific patterns, defining distinct evolutionary routes to resistance.

Altogether, our work delivers the first longitudinal single-cell atlas of metastatic urothelial carcinoma under PD-(L)1 blockade, establishes macrophage polarization as a major driver of resistance, and provides a framework for individualized and dynamically adapted immunotherapeutic strategies.

## Results

### Single-nuclei profiling of metastatic urothelial carcinoma unveils extensive tumor and microenvironmental heterogeneity

A total of 32 patients with metastatic urothelial carcinoma were included in the MATCH-R trial and treated with single-agent PD-1 or PD-L1 inhibitor in the metastatic setting. Median age was 71 (IQR 61-77), and most (66%) had bladder cancer as a primary tumor. Patients were treated either in the first line metastatic setting (28%), second-line (50%), or third-line and more (22%). Most common metastatic sites at immune checkpoint inhibitor initiation were lymph nodes (65%), lung (38%), bone (31%), liver (28%), peritoneal or retroperitoneal soft tissue (28%) (**Supplementary Table S1**). Overall, seven (22%) patients achieved objective response (OR), which translated to prolonged benefit to ICI: with a median follow-up of 26.4 months (95% confidence interval [CI] 16.0-35.4), median progression-free survival was 24.7 months (95%CI 17.8-31.0) in patients who achieved OR versus 1.4 months (95%CI 1.0-1.6) in those who did not, and median overall survival not reached versus 4.2 months (95%CI 2.5-6.2) (**Figure 1a**).

**Figure 1.**
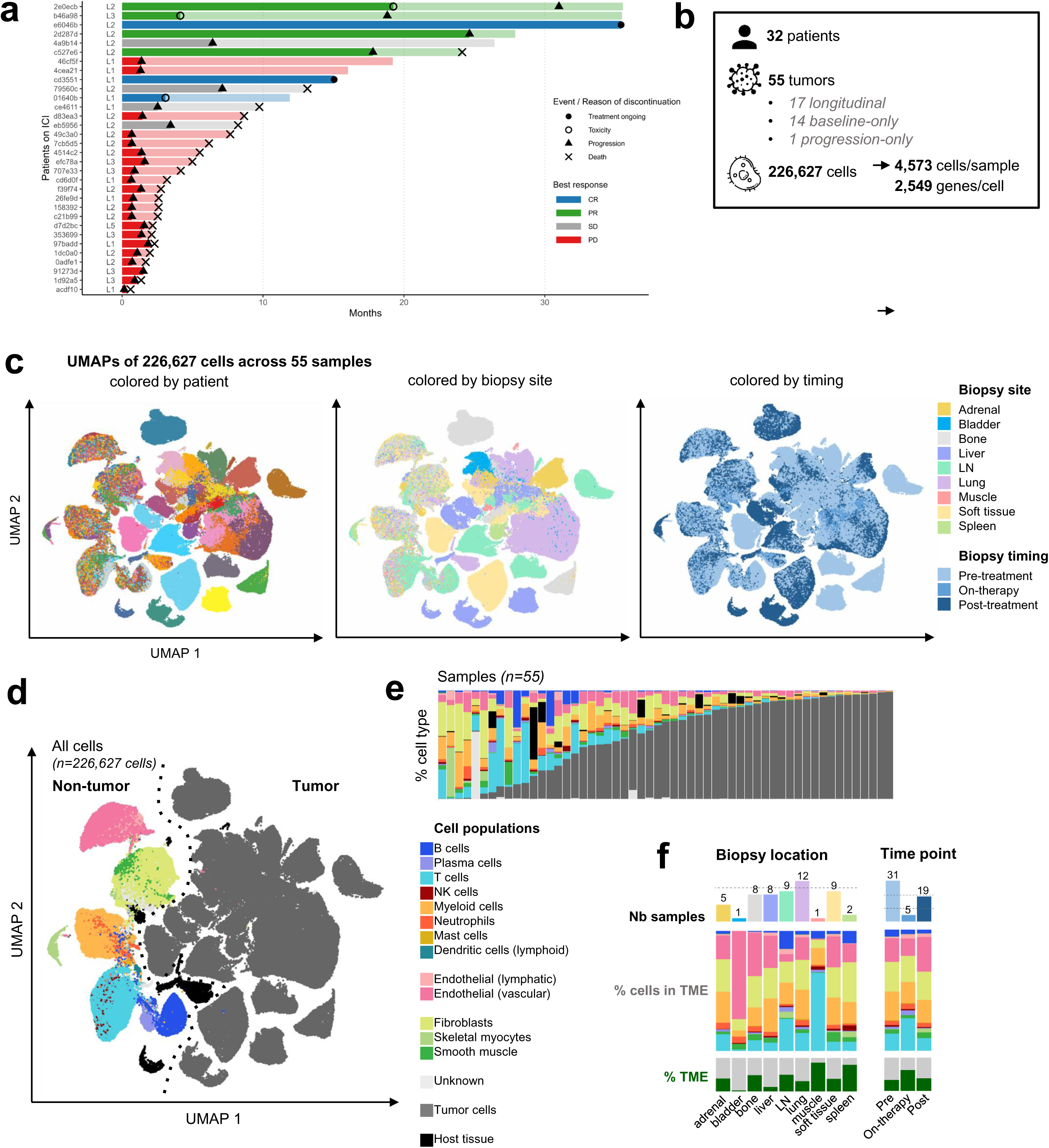
Single-nucleus RNA-seq analysis of 32 patients of the MATCH-R trial. **a.** Swimmer plot depicting individual patient outcomes and biopsy timepoints. **b.** Description of the single-cell MATCH-R cohort. **c.** Uniform manifold approximation and projection (UMAP) of all cells from the MATCH-R cohort (n=226,627 with doublet removal), colored by patient of origin, biopsy site location and treatment timing. **d.** UMAP of cell type annotations, segregated into Tumor and Non-tumor populations. **e.** Overview of cell type abundance across samples. **f.** Abundance of tumor microenvironment (TME) cell populations grouped by biopsyte site location (left) and treatment timing (right).

To allow for longitudinal assessment of the tumor and its microenvironment on therapy, we performed single-nuclei RNA sequencing in matched sequential biopsies at baseline, 3 months after ICI initiation (for patients who achieved disease control), and at disease progression. Biopsies at baseline were performed in 31 patients (97%), including lung (23%), lymph node (19%), soft tissue (19%), liver (13%) or bone (9%) metastases. Seventeen patients (53%) had a biopsy at disease progression, while five (16%) had an interval biopsy on-therapy. Overall, 55 samples were collected. After quality controls (**Supplementary Figure S1**), we obtained single-nuclei transcriptomes from 226,627 cells, with a median number of 4,573 cells per sample, and a median gene coverage of 2,549 (**Figure 1b-c**).

Main immune cell populations were annotated based on the expression of marker genes (see Methods), and tumor cells based on gene expression and virtual copy number profiles (**Figure 1d, Supplementary Figure S2-3**). Tumor cells were the most abundant cell type (mean 59.2% across samples, range 0.2%-98.7%), followed by fibroblasts (mean 11.8%, range 0.1%-68.7%), T cells (mean 8.0%, range 0%-65.6%), macrophages (mean 7.9%, range 0.2%-34.9%), endothelial cells (mean 6.3%, range 0.1%-22.8%) and B cells (mean 2.9%, range 0%-37.7%). Though, a high heterogeneity in immune cell abundance and tumor microenvironment composition was seen across samples and timepoints (**Figure 1e-f**). In particular, biopsies sampled in lymph nodes displayed higher B cell abundance compared to other locations. We also observed an increased abundance of T cells in on-therapy samples (**Figure 1f**).

### Intratumoral coexistence of basal and luminal programs shapes immune infiltration and ICI sensitivity

To explore tumor cell differentiation states, we assessed molecular signatures established from bulk RNA sequencing data^8,14^ in our single-nuclei dataset. We used a pseudobulk approach to classify samples into the consensus MIBC subgroups based on the aggregated single-nuclei transcriptome of their tumor cells. We obtained a high concordance between classifications derived from single-nuclei and matched bulk RNA-seq data (**Supplementary Figure S4**). Yet, individual samples displayed a high heterogeneity regarding tumor cell states, with the coexistence of cells with basal and luminal differentiation (**Figure 2a,b, Supplementary Figure S5**). Principal component analysis revealed two main components significantly associated with the basal, luminal and neuroendocrine differentiation signatures^14^ (**Figure 2a**). Principal component 1 (PC1) was associated with neuroendocrine differentiation (cor = 0.87, P < 2.2E–16), whereas PC2 was positively correlated with the luminal signature and anticorrelated with the basal signature (cor = −0.37, P < 2.2E–16), indicating a continuum between luminal and basal phenotypes.

**Figure 2.**
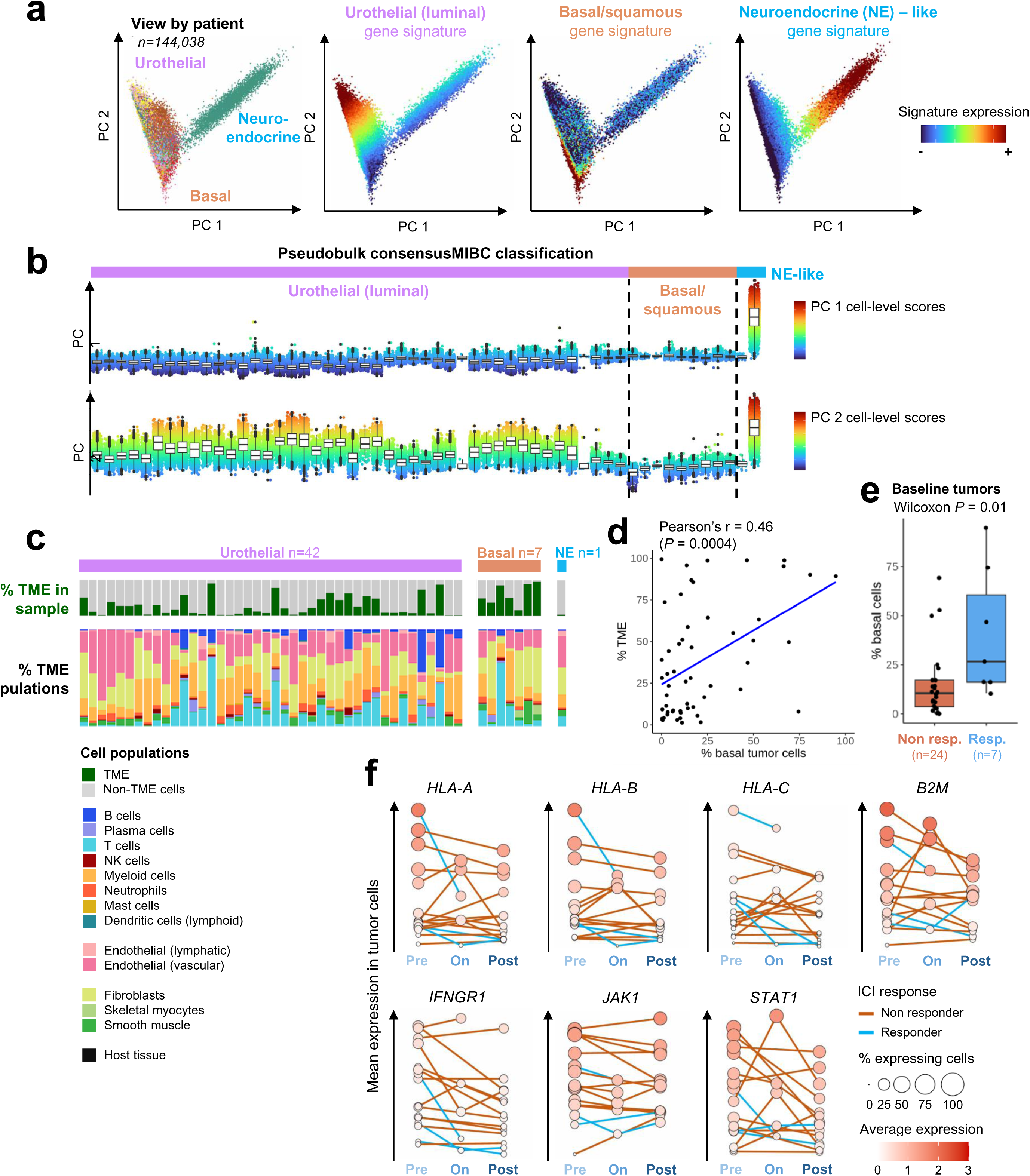
Tumor cell heterogeneity correlates with immune infiltration and ICI response. **a.** Principal component analysis (PCA) of tumor cells (n=144,038), and projection of mean log-normalized expression of urothelial (luminal), basal and neuroendocrine signatures (Biton et al. 2014). **b.** Score distribution of the two first principal components across tumor cells, grouping samples by pseudobulk tumor classification (consensusMIBC). **c.** TME cell type composition according to pseudobulk tumor classification (only samples with ≥ 50 tumor cells were kept). **d.** Correlation between the proportion of basal-like tumor cells and TME abundance across samples. **e.** Proportion of basal tumor cells in baseline tumors from Responder (CR, PR) and Non-responder (PD, SD) patients. **f.** Longitudinal evolution of the expression of antigen presentation and IFN/JAK/STAT pathway genes by tumor cells along treatment time points. Samples from the same patient are linked. The color of the link indicates ICI response.

We then explored whether tumor cell states were associated with specific genomic alterations. We generated whole exome sequencing (WES) data in 32 samples with matched single-cell RNA-seq data. Sixteen potential driver genes were mutated in ≥3 cases including *TP53* (N=26), *KMT2D* (N=18), *TSC1*, *ERBB3* (N=6 each), *KDM6A* (N=5), *ARID1A*, *ERBB2*, *FGFR3*, *HRAS*, *RB1* (N=3 each) (**Supplementary Figure S6a**). We assessed the ability to detect driver alterations in snRNA-seq data, and reached a 74% detection rate for single-nucleotide variants and 83% for oncogene amplifications (**Supplementary Figure S6a,b**). Among putative driver mutations, *KDM6A* was the only one associated with increased PC2 mean (*P* = 0.004) and lower basal cell content (pval *=* 0.01, **Supplementary Figure S6c**). At the intra-tumoral level, we found two genes, *ELF3* (pval *=* 1E-05) and *KDM6A* (pval = 0.01), which mutations were associated with higher cell-wise PC2 values (**Supplementary Figure S6d**).

The relationship between the tumor microenvironment composition and tumor cell states is displayed in **Figure 2c**. Tumors classified as basal/squamous using the pseudobulk classification had increased immune infiltration (**Figure 2c-d** cor=0.46, pval=4E-04), but did not display a significantly different microenvironment composition regarding the main immune cell populations. Despite evidence of higher immune infiltration, the pseudobulk consensus MIBC subgroups did not predict therapy outcome; however, the proportion of basal tumor cells at baseline appeared a significant predictor of response to ICI (pval = 0.01, **Figure 2e**).

### Tumor cells undergo convergent adaptive resistance via loss of antigen presentation and IFN–JAK/STAT signaling

To understand adaptive mechanisms of resistance to ICI related to tumor cells, we assessed gene expression changes between baseline and disease progression. Although no gene was significantly deregulated between pre- and post-treatment in the whole cohort after multiple testing correction, several genes previously associated with immunotherapy response (**Supplementary Table S2**) were significantly down-regulated in post-treatment in individual patients with longitudinal data. We found that genes involved in antigen presentation, including *HLA-A*, *HLA-B*, *HLA-C*, or *B2M* were significantly downregulated in tumor cells at progression in 10/16 patients (**Figure 2f**). Alterations in the JAK/STAT signaling pathway also appeared prominent, with a significant repression of *IFNGR1*, *JAK1* or *STAT1* observed in 11/16 patients with matched samples. In patients who also had tissue sampling during ICI in the context of controlled disease, we observed transient upregulation of *HLA-A/B/C*, *B2M*, *IFNGR1*, and *STAT1* (**Figure 2f**), suggestive of an immune-responsive contexture.

### CD4/CD8 T-cell balance and exhaustion states define response and resistance trajectories

T lymphocytes represented the predominant immune component of the tumor microenvironment in our cohort, with a total of 16,453 cells across the 55 samples (mean = 300 per sample). Following Harmony integration, we identified seven major infiltrating T cell subpopulations: CD4 regulatory cells, CD4 follicular helper cells (naïve, effector, exhausted), and CD8 cytotoxic cells (naïve, effector, exhausted) (**Figure 3a**).

**Figure 3.**
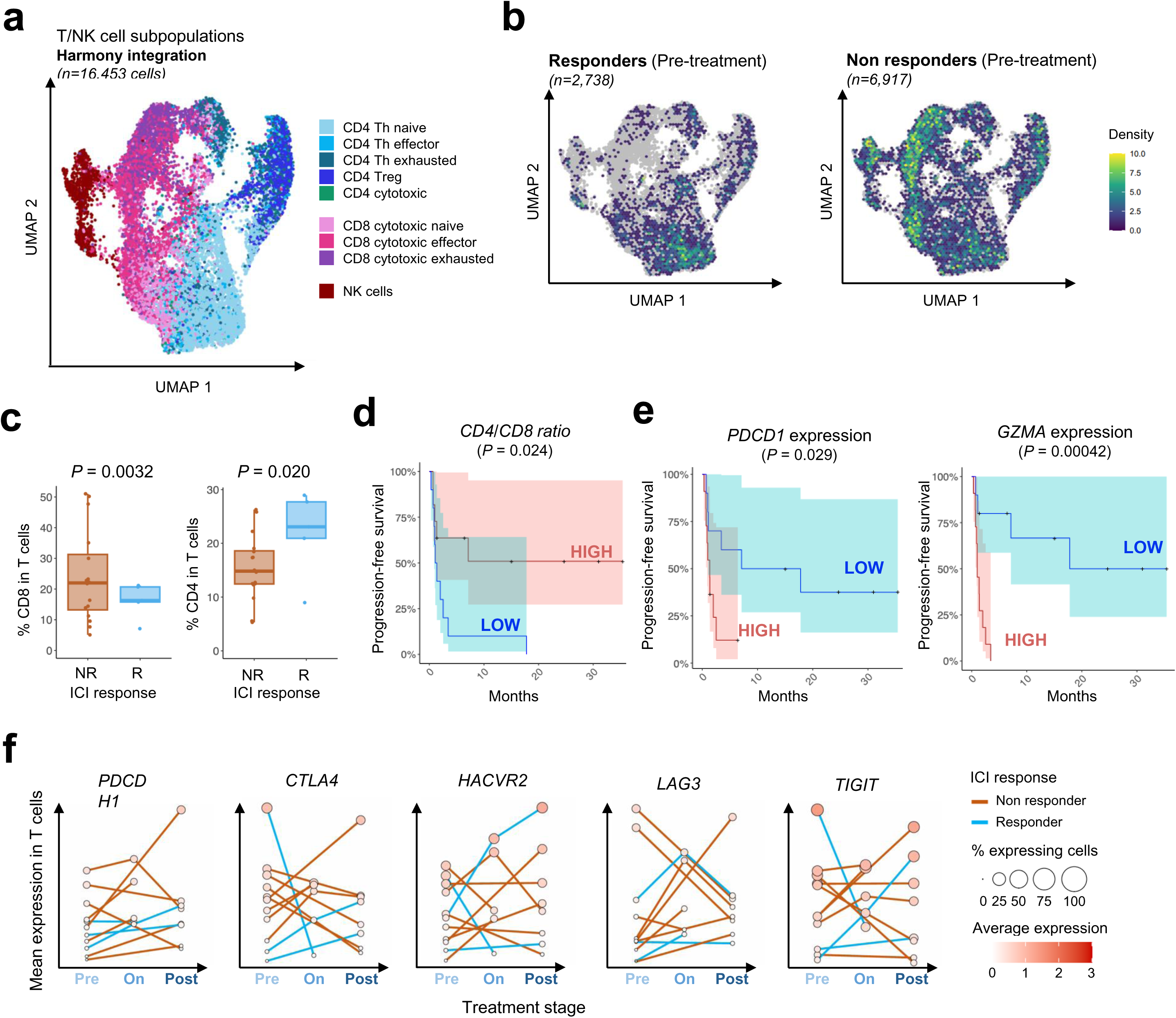
T-cell landscape in metastatic bladder cancer and association with ICI response. **a.** UMAP of T cell and NK cell subpopulations. **b.** UMAP comparison of cell density in baseline tumors of Responder (PR, CR) and Non-responder (PD, SD) patients. **c**. The proportion of CD4 and CD8 T cells are associated with ICI response. **d.** Patients with a higher CD4/CD8 ratio exhibit longer progression-free survival (PFS) after ICI treatment. **e.** Higher *PDCD1* and *GZMA* expression in T cells are associated with shorter PFS after ICI response. High/low expression groups were defined based on median expression across samples. **f.** Longitudinal evolution of the expression of immune checkpoint molecules by T cells during treatment. For each gene, the average log-normalized expression is shown across multiple timepoints (pre-treatment, on-treatment and post-treatment).

The abundance of overall CD8 infiltration was associated with high CD8 effector and CD8 exhausted cell infiltration; both enriched in non-responders (**Figure 3b**). A higher proportion of CD8 T cells among total T cells was associated with poor treatment response (pval = 0.0032, **Figure 3c**), and a lower CD4/CD8 ratio was associated with shorter progression-free survival (pval = 0.024) (**Figure 3d**). At the single-gene level, elevated expression of *PDCD1* and *GZMA* was linked to shorter progression-free survival (**Figure 3e**), suggesting that the CD8 infiltrate in poor responders is enriched for terminally differentiated, exhausted T cells. In contrast, higher overall CD4 T cell infiltration was associated with improved clinical outcomes.

Longitudinal analysis revealed heterogeneous changes in the T cell transcriptome during therapy. Five patients (31%) with sequential T cell single-nuclei data displayed upregulation of at least one immune checkpoint at progression, including *PDCD1*, *CTLA4*, *HAVCR2*, *LAG3* and *TIGIT* (**Figure 3f**).

### Protumoral HES1+ macrophages predict poor response and dominate the evolution of resistance

A total of 12,208 myeloid cells were identified across the 55 samples (mean = 222 per sample). Following Harmony integration, we used the MoMac-VERSE data set comprising a compendium of myeloid cells from multiple single-cell datasets to annotate five macrophage and two monocyte subsets within our clusters of myeloid cells (**Figure 4a**)^15^.

**Figure 4.**
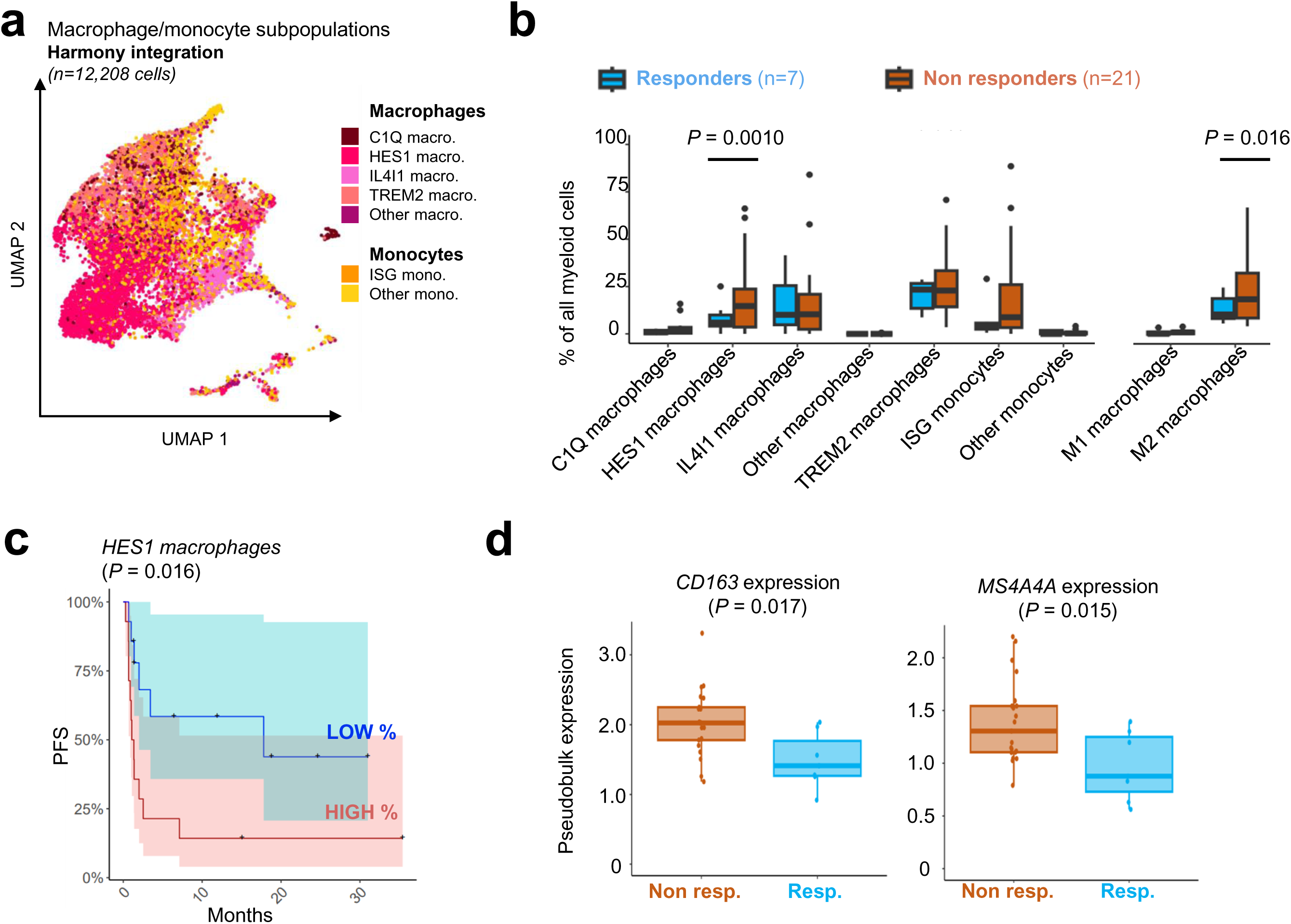
HES1 macrophages predict poor ICI response in mUC. **a.** UMAP of MoMacVerse subpopulations in macrophages and monocytes (n=12,208). **b.** Distribution of macrophage/monocyte subpopulation abundance in baseline tumors from Responder (PR, CR) and Non-responder (PD, SD) patients. **c.** Patients with a higher proportion of HES1 macrophages in myeloid cells at baseline exhibit shorter progression-free survival (PFS) after ICI treatment. **d.** Baseline pseudobulk expression of *CD163* and *MS4A4A* in myeloid cells associates with poor ICI response.

Primary resistance to ICI was associated with higher abundance of pro-tumoral M2 macrophages (*P* = 0.016, GLMM), and in particular HES1 macrophages (*P* = 0.0010, GLMM, **Figure 4b**). A high abundance of HES1 macrophages also predicted shorter PFS (*P* = 0.016, log-rank test, **Figure 4c**). Accordingly, immunosuppressive genes involved in the M2 polarization like *CD163* or *MS4A4A* were upregulated in the macrophages of patients who did not respond to ICI (**Figure 4d**).

In addition, most patients (81%) displayed evidence of a shift towards M2-like macrophage polarization at progression, consisting in increased HES1 macrophage abundance (**Figure 5a**), upregulation of M2-genes such as *CD163* and *VSIG4*, and downregulation of M1 genes such as *CLEC5A* and *HLA-DRA* (**Figure 5b**). This remodeling is illustrated by the UMAPs in **Figure 5c** showing the trajectory of myeloid cells between pre- and post-treatment samples in two patients. Altogether, these data indicated that pro-tumoral macrophages are both implicated in the initial ICI response and in the development of resistance upon treatment.

**Figure 5.**
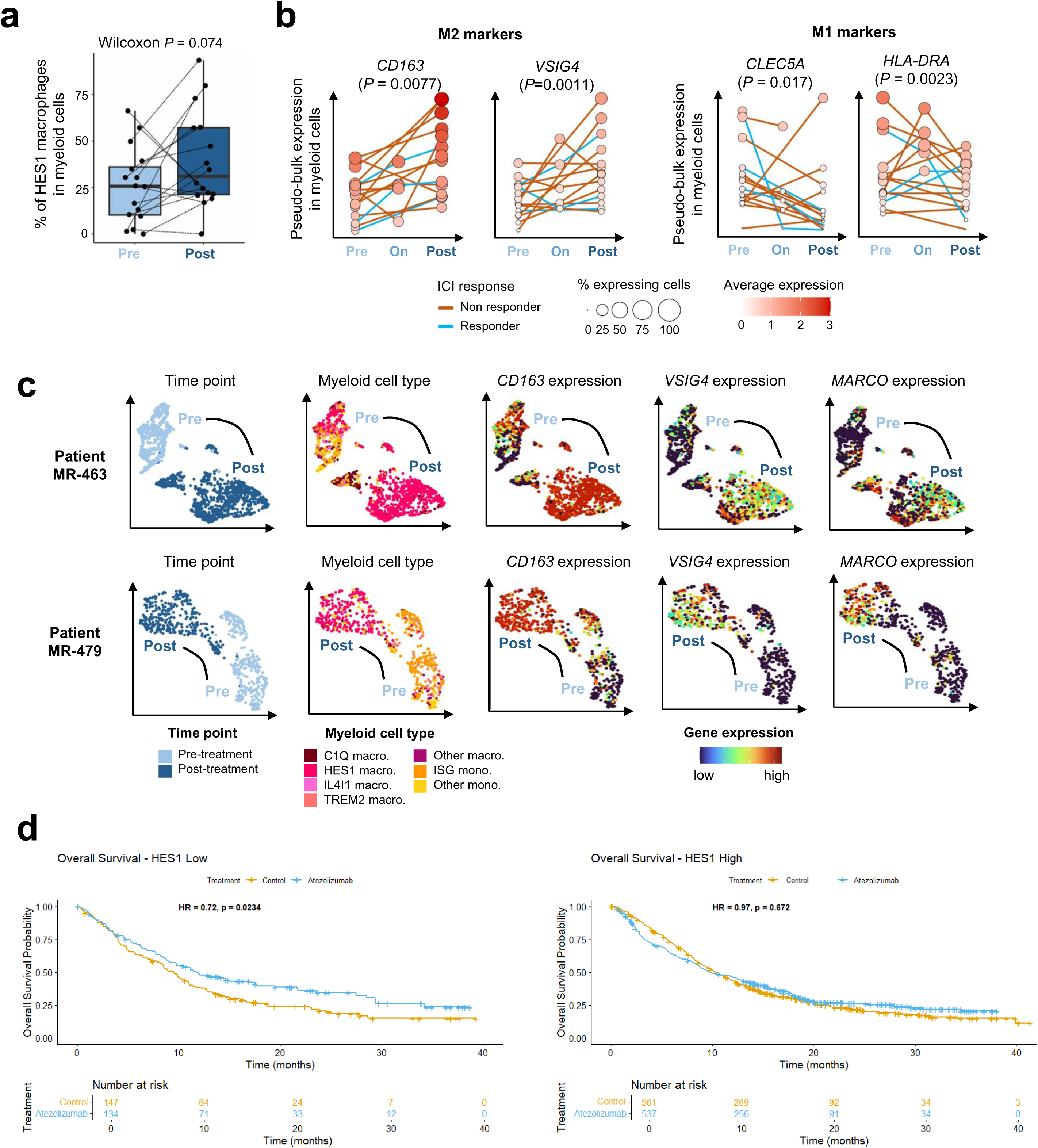
Evidence of protumoral macrophage switch at ICI progression and validation of *HES1* clinical impact in the IMVigor trials a. Evolution the proportion of HES1 macrophages in the myeloid cells of patients with longitudinal samplilng. **b.** Longitudinal evolution of the expression of M1 and M2 markers by myeloid cells along treatment time points. Samples from the same patient are linked. The color of the link indicates ICI response. **c.** Examples of myeloid cell dynamics in two patients with samples pre- and post-ICI treatment. Cells are projected on the UMAPs of myeloid cells and colored from left to right by timing, myeloid cell type, and expression of three markers associated with M2 polarization. **d**. Pooled overall survival analysis of patients with metastatic urothelial carcinoma treated with atezolizumab vs. chemotherapy in the IMVigor 130 and IMVigor211 trials, depending on baseline HES1 expression by bulk RNA sequencing

### Bulk HES1 expression stratifies benefit from PD-L1 blockade across independent clinical trials

Considering that HES1 macrophages were associated with primary and secondary resistance to ICI in our cohort, we evaluated bulk expression of HES1 across publicly available datasets from clinical trials in patients with urothelial cancers, and assessed its impact on outcomes to ICI. Across four trials of atezolizumab in patients with localised or metastatic urothelial cancers, *HES1* expression was significantly increased across stages (metastatic vs. localised) and line of therapy (second-line vs. first-line metastatic setting) **(Supplementary Figure S7).** Pooled overall survival analysis of phase 3 trials evaluating atezolizumab vs. chemotherapy in metastatic urothelial carcinoma showed that only patients with low *HES1* expression derived benefit to atezolizumab versus chemotherapy with an overall survival rate of 48.5% vs. 38.4% at 12 months, and 34.4% vs. 20.8% at 24 months, respectively (HR 0.72, 95% confidence interval [CI] 0.54-0.96). Conversely, patients with high expression of *HES1* did not derive benefit to atezolizumab vs. chemotherapy (HR 0.97, 95%CI 0.84 - 1.12) (**Figure 5d**).

### Patient-specific combinations of tumor and immune adaptations reveal individualized resistance pathways

To recapitulate our findings using more easily accessible assays, we used bulk RNA sequencing to specifically explore genes associated with outcomes through single-nuclei sequencing. We found that candidate genes expression using bulk RNA sequencing was highly driven by the abundance of the tumor microenvironment without discrimination of pro- or antitumor immune features **(Supplementary Figure S8).** Thus, single-nuclei sequencing uniquely allowed the identification of immune features associated with sensitivity to ICI thanks to the depiction of tumor and immune cell subpopulations transcriptomes. Individual patient assessment show specific tumor trajectories involving one or multiple mechanisms of resistance related to tumor or immune cells (**Figure 6**).

**Figure 6.**
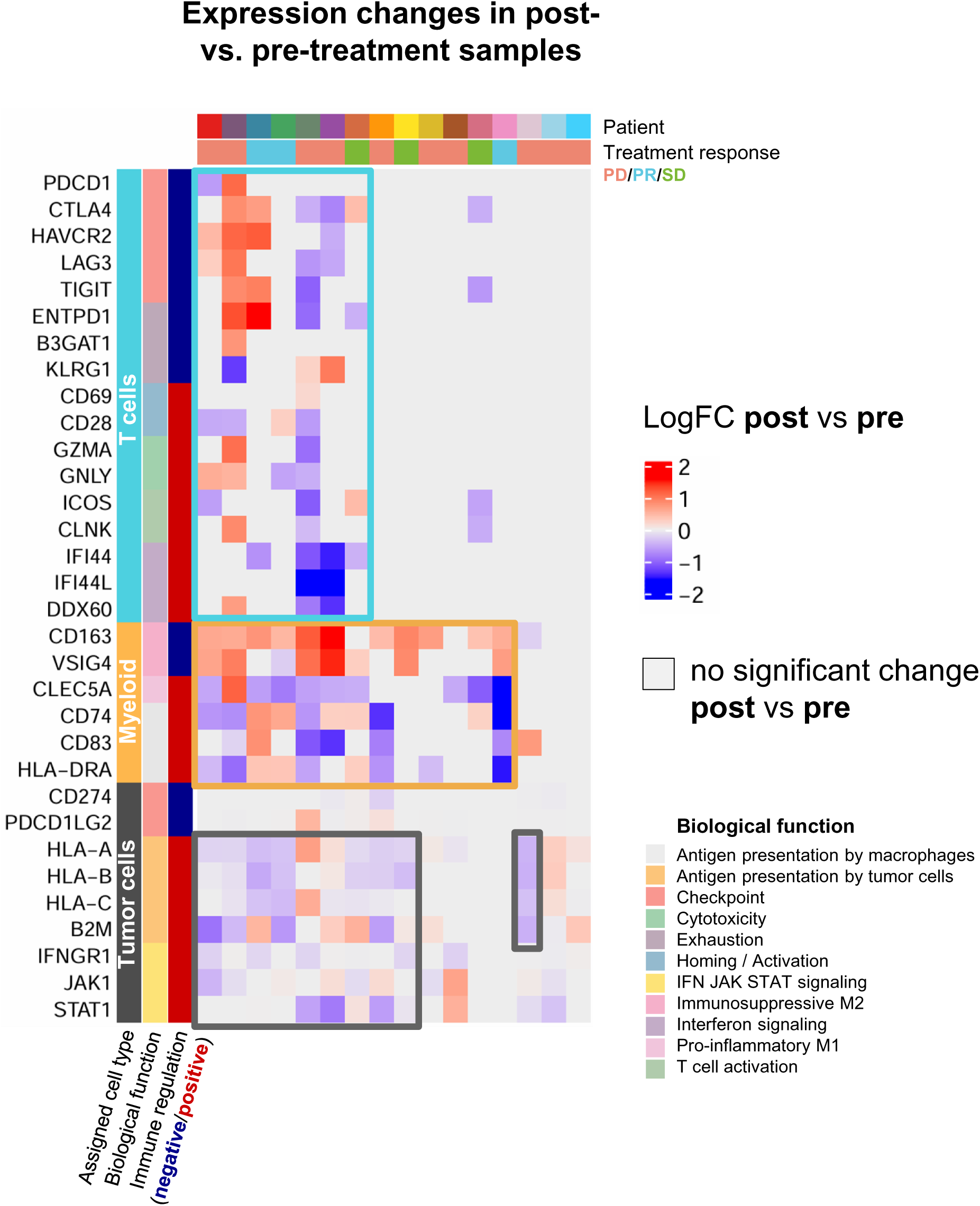
Overview of individual patient trajectories involving tumor and immune cell reprogramming after immune checkpoint inhibition. Each column corresponds to one patient with longitudinal sampling, and each line to a gene, grouped by cell type of interest. The heatmap indicates with a color code whether each gene is significantly up-(red) or down-regulated (blue) in the post-treatment sample compared to the pre-treatment sample. Absence of differential expression is shown in grey.

## Discussion

Using longitudinal single-nuclei sequencing of metastatic biopsies, this study delineates dynamic tumor-immune trajectories underlying response and resistance to PD-(L)1 blockade in metastatic urothelial carcinoma. By resolving cell-state-specific programs across tumor, lymphoid, and myeloid compartments, we identify macrophage-driven tumor-immune reprogramming as a dominant and convergent mechanism of both primary and acquired resistance to immune checkpoint inhibition.

A major strength of this work lies in its focus on metastatic lesions sampled longitudinally, a setting in which resistance to immune checkpoint inhibitors ultimately emerges and therapeutic decisions are made. While previous studies have largely relied on bulk transcriptomics or primary tumor samples, our approach captures the evolving cellular ecosystems of metastatic disease at single-cell resolution, revealing resistance mechanisms that are both dynamic and patient specific.

First, we observed that tumor cell plasticity modulates, but does not dominate, resistance trajectories. At the tumor cell level, we found extensive intratumoral heterogeneity, with basal and luminal differentiation states coexisting within individual metastatic lesions. This plasticity challenges the predictive value of molecular subtyping derived from bulk analyses and suggests that tumor cell state composition, rather than subtype assignment, more accurately reflects immunotherapy sensitivity. Indeed, the impact of tumor transcriptome on outcomes for patients treated with ICI has been controversial in studies using bulk transcriptome datasets. In localized trials evaluating PD-(L)1 inhibition as neoadjuvant therapy, the PURE01 trial showed that patients with basal phenotypes had highest response rates, a finding that was not confirmed in the ABACUS trial^9^. In the metastatic setting, the atlas published by Loriot et al. showed that bulk transcriptome was not indicative of immune infiltration^16^. Our study showed that a higher proportion of basal-like tumor cells was associated with increased immune infiltration and improved response to PD-(L)1^8,13^. Longitudinal analyses further reveal adaptive tumor-intrinsic mechanisms at progression, including downregulation of antigen presentation machinery and interferon signaling pathways. These changes likely contribute to immune escape in a subset of patients, echoing resistance mechanisms described across multiple cancer types. However, the heterogeneity and incomplete penetrance of these alterations suggest that tumor-intrinsic reprogramming alone is not enough to explain resistance in most patients.

Lymphocyte infiltration has been usually positively associated with outcomes, with B and T cell lineages repeatedly associated with improved response to ICI^7,9,10^. Though, it has been shown that many patients with high lymphocyte infiltration may not derive benefit from ICI. In the study from Robertson et al. based on localized tumors treated with ICI, patients who were non-responders despite high lymphocyte infiltration harbored high checkpoint expression^9^. In the lymphoid compartment, we found that resistance to PD-(L)1 blockade is characterized not by absence of immune infiltration, but the predominance of dysfunctional immune states. Our work confirms that high checkpoint expression on CD8 lymphocytes is reflective of an exhausted immune infiltrate, which is both a primary and adaptive mechanisms of resistance to ICI. In contrast, greater CD4 T cell abundance correlated with improves outcomes supporting a model in which coordinated helper T cell responses are required for durable antitumor response. Overall CD4 infiltration was associated with better outcomes, although individual CD4 cell subpopulations were not strong predictors of ICI activity in our dataset. The important role of CD4 in response to ICI has been suggested by Goubet et al.^7^, who found that T follicular helpers were predictors of benefit in localized ICI-treated patients, and that CD4 T cells had highest binding potential to pembrolizumab. B cells, and notably antibody-secreting cells, has also been recently proposed as important mediators of immune response^7^. Though, B cell infiltration in our study was mostly restricted to patients with lymph node metastases, a metastatic location usually associated with better outcomes on ICI^17^. However, given the physiological abundance of B cells in lymph nodes, broader conclusions about B cell subtyping in this metastatic setting is not currently possible. Small biopsy fragments in our metastatic study may have also limited the detection of tertiary lymphoid structures, key actors of the coordination of the immune response and host of B and T follicular helper interactions^7,18^, hence impacting our ability to fully capture B cell and CD4 subtype influence on ICI response. Longitudinal upregulation of inhibitory receptors on T cells at progression further underscores immune exhaustion as an adaptive resistance mechanism. However, as with tumor-intrinsic changes, these lymphoid alterations were variable across patients and frequently coexisted with other resistance programs, indicating that T cell dysfunction is part of a broader, multidimensional resistance landscape.

In contrast to the heterogeneous contributions of tumor and lymphoid compartments, myeloid cell trajectories revealed a strikingly consistent pattern. Both primary resistance and disease progression were characterized by enrichment of protumoral macrophage states, particularly a HES1 expressing macrophage population associated with immunosuppressive gene expression and worst clinical outcomes. Evidence of protumoral polarization, notably through HES1 and TREM2 subpopulations abundance, drove primary and secondary resistance to ICI, while M2-like and M1-like genes expression had opposite trajectories from treatment-start to treatment-resistance. This convergence suggests that macrophage reprogramming represents a dominant resistance axis. So far, very limited evidence was available to suggest a leading role for tumor infiltrating macrophages in urothelial cancers. Data from the IMvigor210 trial corroborated that M1-like macrophage signatures could be associated with improved outcomes in patients treated with atezolizumab. A broader bulk RNA sequencing dataset in metastatic patients treated with atezolizumab associated a “myeloid” signature with immune-responsive tumors, but also with higher expression of immune checkpoints^10^. This divergence from our findings may stem from the absence of clear pro- or anti-tumoral feature set in the “myeloid” signature explored in this study, as well as the fact that most samples stemmed from primary tumors that may not reflect metastatic states. This limited granularity may also explain that the “myeloid” signature in this trial harbored conflicting predictive ability depending on tumor subtypes, demonstrating the importance of accurate single-cell state determination. Together, these data position macrophage-driven immunosuppression as a central determinant of ICI failure.

Our findings have important implications for biomarker development. Bulk transcriptomic signatures largely reflect overall immune abundance and fail to discriminate protumoral from antitumoral immune states, limiting their predictive utility. In contrast, single-cell resolved macrophages states, such as HES1 expressing populations capture functionally relevant resistance programs and may inform patient stratification for immunotherapy. Therapeutically, our results provide a strong rationale for targeting macrophage-driven immunosuppression in combination with PD-(L)1 inhibitors. Strategies aimed at reprogramming tumor-associated macrophages or disrupting protumoral myeloid signaling pathways could restore immune sensitivity in both primary and acquired resistance settings.

This study has limitations inherent to its design. The cohort size is modest, reflecting the technical and logistical challenges of longitudinal metastatic biopsies and sampling of small core biopsies may underrepresent spatial immune structures such as tertiary lymphoid structures. Limited tumor material availability could also explain the low number of dendritic cells in our samples, although those have been previously reported to be present in immune-responsive tumor subtypes^10^. Assessment of the greater stromal architecture may also be hampered by the small sampling size. This could also explain why cancer associated fibroblasts did not emerge as a potential predictor of outcomes in our study, while those have been previously associated with stromal, immune-resistant tumors^10,19^. Nevertheless, the depth of longitudinal sampling and the consistency of macrophage-driven resistance across patients and external datasets mitigate these limitations. Future studies integrating spatial transcriptomics, functional perturbation models and prospective clinical validation will be required to fully elucidate the regulatory network governing macrophage reprogramming and to translate these insights inti therapeutic approaches.

In conclusion, longitudinal single-cell profiling of metastatic urothelial carcinoma reveals that resistance to PD-(L)1 blockade converges on macrophage driven tumor-immune reprogramming. While tumor cell plasticity and T cell exhaustion contribute to resistance in selected contexts, protumoral macrophage polarization emerges as a dominant and recurrent mechanism underlying both primary and acquired immune checkpoint inhibitor failure. These data support the development of macrophage-targeted combination strategies in metastatic urothelial carcinoma.

## Material and methods

### Single-nuclei RNA sequencing

Cryopreserved cells were thawed and nucleus permeabilization was performed following recommendations from 10X Genomics. Single-nucleus 5’ gene expression libraries were generated using 10X Genomics single cell gene expression 5’ v1.1 kit, following the protocol provided by the manufacturer. A total of 10,000 nuclei per sample were loaded in each lane of the Chromium Single Cell Controller, unless fewer nuclei were recovered after dissociation, in which case they were all loaded. Libraries were sequenced on a NovaSeq 6000 to a target depth of 150 million reads (Paired-end, 28bp Read 1, 90bp Read 2), and topped-up accordingly in order to reach sequencing saturation for each library.

### Single-nuclei RNA-seq processing

We used a custom workflow to process raw FASTQ files. The UMI counts matrices were generated using STAR^20^ alignment and count (STARsolo) with the human genome reference (GrCh38/hg38) and 10x barcode whitelists. We used CellBender^21^ to remove counts from ambient RNA molecules and random barcode swapping with recommended settings. We used DoubletCollection^22^ to detect doublet cells with *doubletCells*, *cxds, bcds*, *hybrid, scDblFinder, Scrublet, DoubletDetection* and *DoubletFinder* algorithms. Each cell detected as doublet by ≥3 methods was removed. We retained only high-quality cells (>500 genes detected and <10% of mitochondrial RNA reads). The samples were aggregated into one matrix and the cohort matrix was normalized using monocle3^23^ normalization method by size factor. We then performed dimensionality reduction using principal component analysis (PCA) (producing 20 principal components) and UMAP (Uniform Manifold Approximation and Projection) on the first 20 principal components. Last, we applied PhenoGraph clustering (with Leiden algorithm) with a resolution of 0.3 on the 20 PCA components.

### Annotation of broad cell types

We annotated cells by projecting the expression of established marker genes on the PhenoGraph clusters obtained during the previous step. Each cluster was assigned to the cell type with highest marker expression, and doublets were removed. This annotation step enabled to identify potential bladder tumor cells as well as cells from the tumor microenvironment (TME: fibroblasts, T cells, macrophages, endothelial cells, B cells, neutrophils) and from the host tissue of the metastasis biopsy (lung, liver, adrenal). We ran infercnvpy^24^ to compute a CNV score per cell, taking TME populations as reference cells. We then checked that all epithelial cells displayed higher CNV scores compared to the reference cells and labeled these as “tumor cells”. To refine the annotation of TME populations, we performed a second run of dimensionality reduction and clustering exclusively on the TME cells (n=77,014) Using established marker genes, we could annotate finer populations such as vascular cells, smooth muscle cells, plasma cells, skeletal myocytes, dendritic cells or mast cells.

### Detailed annotation of T cells

To annotate in-depth the functional subsets of T cells, we used the Seurat package^25^ to process the corresponding datasets as follows: log-normalization (NormalizeData), identification of the most variable genes (FindVariableFeatures with vst), scaling (ScaleData), dimensionality reduction with PCA (RunPCA with 30 components), identification of transcriptomic clusters (FindNeighbors and FindClusters with a resolution of 0.5 for T cells and 1 for myeloid cells) and UMAP visualization (runUMAP). We used the harmony^26^ package (RunHarmony with default parameters) to integrate cells across multiple biopsy sites. We then used the *UCell* ^27^ package to compute cell-level scores associated to CD4 T cell markers (*CD4*, *IL7R*, *FOXP3*, *IKZF2*, *CXCR3*, *CCR6*, *IL2RA*, *LEF1*) and CD8 T cell markers (*CD8A*, *CD8B*, *GZMB*, *PRF1*, *KLRD1*, *NKG7*, *IFNG*, *CCL4*, *EOMES*). T cell Seurat clusters were annotated as CD4/CD8 based on their average UCell scores, and cells clusters with high expression of NK cell markers (*NCAM1*, *KLRF1*, *KLRC1*, *KLRD1* and *FCGR3A*) were labeled “NK cells”. We then isolated CD4 and CD8 T cells and computed UCell scores per cell using five CD4-related signatures (naïve, effector, exhausted, cytotoxic, regulatory) and three CD8-related signatures (naïve, effector, exhausted) extracted from a published pan-cancer T cell atlas^28^. We kept genes with sufficient expression in the dataset (at least 1 count in 1% cells) and removed genes overlapping multiple signatures. UCell scores were scaled between 0 and 1 for each signature. Cells with all scores lower than 0.1 were labeled “Unknown”, while remaining cells were labeled according to the state with the highest UCell score.

### Detailed annotation of myeloid cells

We annotated myeloid cells into the functional subsets identified in the MoMacVerse meta-analysis ^15^. After integrating and clustering myeloid cells from all samples using the same approach as for T cells, we used Seurat *FindAllMarkers* function to identify markers of each myeloid subset defined in the MoMacVerse data set. For each subset, we used as markers the 50 genes with the highest fold-change compared to other subsets, within significant genes (p-value < 0.05 and fold-change > 1.5). We kept genes with sufficient expression in our data set (at least 1 count in 1% of cells) and removed genes overlapping mutliple signatures. Signatures comprising ≤5 genes were discarded. UCell scores were scaled between 0 and 1 for each signature. Cells with all scores lower than 0.1 were labeled “Unknown”, while other cells were labeled according to the myeloid subset with the highest UCell score. Functionally similar subsets of myeloid cells were merged, leading to a final annotation into five macrophage subsets (HES1, TREM2, C1Q, IL4I1 and others) and two monocyte subsets (ISG and others).

### Bladder cancer cell states

To explore the heterogeneity of bladder cancer cells, we conducted a PCA analysis with the Seurat package (20 components) on 103 genes previously associated with urothelial, basal and neuroendocrine differentiation signatures in bladder cancer^29^. Projected tumor cells on the two first components revealed a clear separation of neuroendocrine cells but a gradient between the most basal and luminal differentiation. We then computed the log-normalized mean expression of each set of marker genes in each cell. The proportion of basal cells in each sample was estimated as the proportion of cells with a log-normalized expression of the basal signature ≥ 0.1. We applied consensusMIBC^29^ on pseudo-bulk RNA profiles generated by pooling tumor cell profiles to predict the molecular class of each sample and we compared the results to consensusMIBC annotations based on matched bulk RNA-seq data.

### Association with clinical response

To identify predictors of ICI response, we compared cell type proportions and expression of individual genes between baseline samples of responders and non-responders. We used quasi-binomial generalized linear mixed models (GLMM function from the *lme4* ^30^ R package with logit link, using the biopsy site as random effect) to compare the abundance of immune cell subsets (or cells positive for a given functional signature) between responders and non-responders. To evaluate the expression of individual genes within cell types of interest (e.g. *PDCD1* in T cells), we computed pseudobulk expression levels and we used limma^31^ package to identify differentially expressed genes between responders and non-responders. We used log-rank tests (*survival* ^32^ R package) to compare the progression-free survival (PFS) and overall survival (OS) between patients with low/high scores for each feature, split by the median.

### Longitudinal evolution upon treatment

To identify longitudinal gene expression changes in each cell type, we performed differential expression analyses between pre- and post-treatment samples using Seurat *FindMarkers* with ‘test.use=MAST’ and ‘latent.var=patient’. For genes of interest, e.g. genes known to be associated with ICI response, and genes significantly deregulated between pre- and post-treatment samples, we generated visualizations showing the proportion of cells expressing each gene and the mean expression level in matched pre/post pairs.

### Mutation analyses

To investigate mutation detection at the single-cell level, we ran the scReadCounts ^33^ tool on the scRNA-seq BAM alignment files of 32 bladder tumors with paired bulk whole exome sequencing (WES) data. To identify somatic mutations from WES data, reads were aligned to the human genome build hg38/GRCh38.p7 using the Burrows-Wheeler Aligner (BWA) tool^34^ (v0.7.17-r1188). Duplicated reads were removed using Sambamba^35^. Somatic single nucleotide variants (SNVs) and small insertions/deletions (indels) were detected using MuTect2^36^ tool (2.0, --max_alt_alleles_in_normal_count=2; --max_alt_allele_in_normal_fraction=0.04) and annotated using Ensembl’s Variant Effect Predictor^37^ (VEP, release 101). We next screened identified somatic mutations in snRNA-seq data. For each sample, scReadCounts computed the number of reference (REF) or alternate/mutated (ALT) reads for each somatic coding SNV detected in matched WES data. Cells were considered mutated for a position if they displayed >= 1 mutated (ALT) read for the position. Mutated (ALT) and reference (REF) reads were aggregated by cell population (Tumor or TME), and mutations were labeled as follows: “Detected” if they displayed >= 3 mutated (ALT) reads in the tumor population, “Not detected” if they displayed < 3 mutated (ALT) reads and >= 10 total reads in the tumor population, “Not covered” if they displayed < 10 total reads in the tumor population.

### Gene Expression Analysis and Survival Correlation

Bulk RNA-sequencing and clinical data were obtained from the European Genome-phenome Archive (EGA) for four clinical trials evaluating atezolizumab in bladder cancer (EGAS50000000497). Gene expression data were provided as log-transformed transcripts per million (logTPM) values. HES1 expression was analyzed across all studies using the Kruskal-Wallis test to assess inter-study distribution differences. For survival analysis, the optimal cutpoint for HES1 expression was determined using the maximally selected rank statistics method (surv_cutpoint function from the survminer^38^ R package) in the atezolizumab treatment arm of metastatic patients. Patients were classified as HES1-high or HES1-low based on this cutpoint. Overall survival (OS) was estimated using the Kaplan-Meier method, and differences between groups were assessed using the log-rank test. To assess outcomes according to HES1 in the metastatic setting, data from the two randomised phase 3 studies IMVigor130 and IMVigor211 were combined, excluding patients treated with the atezolizumab-chemotherapy combination arm to focus on monotherapy comparisons (atezolizumab vs. chemotherapy alone). Cox proportional hazards regression was performed within each HES1 expression subgroup to evaluate the treatment effect (atezolizumab vs. control) and to test for treatment-biomarker interaction. Median follow-up was calculated using the reverse Kaplan-Meier method. All statistical analyses were performed using R version 4.2.0 with the survival, survminer, dplyr, and ggplot2 packages. A two-sided p-value < 0.05 was considered statistically significant.

## Acknowledgements

This work, as part of IHU PRISM National PRecISion Medicine Center in Oncology, received state funding managed by the French National Research Agency under the France 2030 program (grant no. ANR-23-IAHU-0002).

Ronan Flippot was funded by the ARC foundation for cancer research and the BMS foundation.

---

**Supplementary Table S2.** Candidate genes involved in immunotherapy.

**Supplementary Figure S1.** Description of quality control metrics per sample (from top to bottom: number of cells in sample, total counts/cell in sample, gene coverage/cell in sample, mitochondrial RNA percentage/cell in sample). Red full lines highlight median values across the cohort for each metric.

**Supplementary Figure S2.** Annotations of subpopulations in TME cells (top) and expression of correpsonding markers (bottom).

**Supplementary Figure S3.** UMAP of Copy Number Variation (CNV) scores generated by inferCNVpy projected in tumor cells and TME cells.

**Supplementary Figure S4.** Consensus classification (Kamoun et al. 2020) of samples according to pseudobulk single-nuclei sequencing and bulk transcriptome (N=30).

**Supplementary Figure S5.** Heterogeneity of single-nuclei tumor cell transcriptome by sample. Samples are ordered by mean expression difference between basal (n=143 genes) and urothelial (luminal, n=194 genes) differentiation signatures.

**Supplementary Figure S6.** Single-nuclei detection of driver alterations. **a.** Single-cell detection of driver mutations identified by whole exome sequencing. **b.** Single-cell detection of high-level amplifications identified by whole exome sequencing. **c.** *KDM6A*-mutated samples display significantly lower basal cell content and higher PC2 values. **d.** Cells in which *ELF3* and *KDM6A* mutations were detected display significantly higher PC2 values.

**Supplementary Figure S7.** Expression of HES1 across pivotal trials of atezolizumab in urothelial cancers across stages and lines of therapy.

**Supplementary Figure S8.** (Top) Bulk RNA sequencing of the MATCH-R baseline cohort displaying inferred MCP-Counter scored immune cell types and (bottom) expression of immune-related genes according to response to therapy

**Supplementary Table S1.** Baseline characteristics of the population.

|  |  | Study population<br>N=32 (%) |
| --- | --- | --- |
| **Age** |  |  |
| median, years (IQR) |  | 71 (61-77) |
| **Sex** |  |  |
| Male |  | 24 (75) |
| **Primary tumor** |  |  |
| *Bladder* |  | 21 (66) |
| *Upper Tract* |  | 9 (28) |
| *Urethra* |  | 1 (3) |
| *Multifocal* |  | 1 (3) |
| **Metastatic sites** |  |  |
| *Lymph Nodes* |  | 21 (65) |
| *Lung* |  | 12 (38) |
| *Bone* |  | 9 (28) |
| *Liver* |  | 9 (28) |
| *Soft tissue* |  | 10 (31) |
| *Adrenal* |  | 3 (9) |
| **Line of therapy** |  |  |
| 1 |  | 9 (28) |
| 2 |  | 16 (50) |
| 3+ |  | 7 (22) |
| **Immune checkpoint inhibitor** |  |  |
| *Pembrolizumab* |  | 30 (94) |
| *Durvalumab* |  | 1 (3) |
| *Avelumab* |  | 1 (3) |

## References.

1. Bray, F. et al. Global cancer statistics 2022: GLOBOCAN estimates of incidence and mortality worldwide for 36 cancers in 185 countries. CA Cancer J Clin 74, 229–263 (2024).

2. van der Heijden, M. S., et al. Nivolumab plus Gemcitabine–Cisplatin in Advanced Urothelial Carcinoma. New England Journal of Medicine 389, 1778–1789 (2023).

3. Powles, T., et al. Avelumab Maintenance Therapy for Advanced or Metastatic Urothelial Carcinoma. N Engl J Med 383, 1218–1230 (2020).

4. Bellmunt, J., et al. Pembrolizumab as Second-Line Therapy for Advanced Urothelial Carcinoma. New England Journal of Medicine 376, 1015–1026 (2017).

5. Balar, A. V., et al. Efficacy and safety of pembrolizumab in metastatic urothelial carcinoma: results from KEYNOTE-045 and KEYNOTE-052 after up to 5 years of follow-up☆. Annals of Oncology 34, 289–299 (2023).

6. Powles Thomas et al. Enfortumab Vedotin and Pembrolizumab in Untreated Advanced Urothelial Cancer. New England Journal of Medicine 390, 875–888 (2024).

7. Goubet, A.-G., et al. Escherichia coli-Specific CXCL13-Producing TFH Are Associated with Clinical Efficacy of Neoadjuvant PD-1 Blockade against Muscle-Invasive Bladder Cancer. Cancer Discov 12, 2280–2307 (2022).

8. Kamoun, A., et al. A Consensus Molecular Classification of Muscle-invasive Bladder Cancer. European Urology 77, 420–433 (2020).

9. Robertson, A. G., et al. Expression-based subtypes define pathologic response to neoadjuvant immune-checkpoint inhibitors in muscle-invasive bladder cancer. Nat Commun 14, 2126 (2023).

10. Hamidi, H., et al. Molecular heterogeneity in urothelial carcinoma and determinants of clinical benefit to PD-L1 blockade. Cancer Cell 42, 2098–2112.e4 (2024).

11. Gao, J., et al. Neoadjuvant PD-L1 plus CTLA-4 blockade in patients with cisplatin-ineligible operable high-risk urothelial carcinoma. Nat Med 26, 1845–1851 (2020).

12. van Dijk, N., et al. Preoperative ipilimumab plus nivolumab in locoregionally advanced urothelial cancer: the NABUCCO trial. Nat Med 26, 1839–1844 (2020).

13. Necchi, A., et al. Impact of Molecular Subtyping and Immune Infiltration on Pathological Response and Outcome Following Neoadjuvant Pembrolizumab in Muscle-invasive Bladder Cancer. Eur Urol 77, 701–710 (2020).

14. Biton, A., et al. Independent component analysis uncovers the landscape of the bladder tumor transcriptome and reveals insights into luminal and basal subtypes. Cell Rep 9, 1235–1245 (2014).

15. Mulder, K., et al. Cross-tissue single-cell landscape of human monocytes and macrophages in health and disease. Immunity 54, 1883–1900.e5 (2021).

16. Loriot, Y., et al. The genomic and transcriptomic landscape of metastastic urothelial cancer. Nat Commun 15, 8603 (2024).

17. Bellmunt, J., et al. Avelumab first-line maintenance (1LM) for advanced urothelial carcinoma (aUC): Long-term outcomes from JAVELIN Bladder 100 in patients (pts) with low tumor burden. JCO 42, 4566–4566 (2024).

18. Flippot, R., et al. B cells and the coordination of immune checkpoint inhibitor response in patients with solid tumors. J Immunother Cancer 12, e008636 (2024).

19. Chen, Z., et al. Single-cell RNA sequencing highlights the role of inflammatory cancer-associated fibroblasts in bladder urothelial carcinoma. Nat Commun 11, 5077 (2020).

20. Dobin, A., et al. STAR: ultrafast universal RNA-seq aligner. Bioinformatics 29, 15–21 (2013).

21. Fleming, S. J., et al. Unsupervised removal of systematic background noise from droplet-based single-cell experiments using CellBender. Nat Methods 20, 1323–1335 (2023).

22. Xi, N. M. & Li, J. J. Protocol for executing and benchmarking eight computational doublet-detection methods in single-cell RNA sequencing data analysis. STAR Protoc 2, 100699 (2021).

23. Trapnell, C., et al. The dynamics and regulators of cell fate decisions are revealed by pseudotemporal ordering of single cells. Nat Biotechnol 32, 381–386 (2014).

24. infercnvpy: Scanpy plugin to infer copy number variation (CNV) from single-cell transcriptomics data. Project name not set https://infercnvpy.readthedocs.io/en/latest/index.html.

25. Hao, Y., et al. Integrated analysis of multimodal single-cell data. Cell 184, 3573–3587.e29 (2021).

26. Korsunsky, I., et al. Fast, sensitive and accurate integration of single-cell data with Harmony. Nat Methods 16, 1289–1296 (2019).

27. Andreatta, M. & Carmona, S. J. UCell: Robust and scalable single-cell gene signature scoring. Computational and Structural Biotechnology Journal 19, 3796–3798 (2021).

28. Chu, Y., et al. Pan-cancer T cell atlas links a cellular stress response state to immunotherapy resistance. Nat Med 29, 1550–1562 (2023).

29. Kamoun, A., et al. A Consensus Molecular Classification of Muscle-invasive Bladder Cancer. Eur Urol 77, 420–433 (2020).

30. Bates, D., Mächler, M., Bolker, B. & Walker, S. Fitting Linear Mixed-Effects Models Using lme4. Journal of Statistical Software 67, 1–48 (2015).

31. Ritchie, M. E., et al. limma powers differential expression analyses for RNA-sequencing and microarray studies. Nucleic Acids Res 43, e47 (2015).

32. Therneau, T. A package for survival analysis in R.

33. Prashant, N. M., et al. SCReadCounts: estimation of cell-level SNVs expression from scRNA-seq data. BMC Genomics 22, 689 (2021).

34. Li, H. & Durbin, R. Fast and accurate short read alignment with Burrows-Wheeler transform. Bioinformatics 25, 1754–1760 (2009).

35. Tarasov, A., Vilella, A. J., Cuppen, E., Nijman, I. J. & Prins, P. Sambamba: fast processing of NGS alignment formats. Bioinformatics 31, 2032–2034 (2015).

36. Cibulskis, K., et al. Sensitive detection of somatic point mutations in impure and heterogeneous cancer samples. Nat Biotechnol 31, 213–219 (2013).

37. McLaren, W., et al. The Ensembl Variant Effect Predictor. Genome Biol 17, 122 (2016).

38. Kassambara, A. kassambara/survminer. (2025).

