## Supplemental_Figures for "Single-cell trajectories in metastatic urothelial carcinoma reveal tumor–immune reprogramming and macrophage-driven resistance to PD-(L)1 blockade"

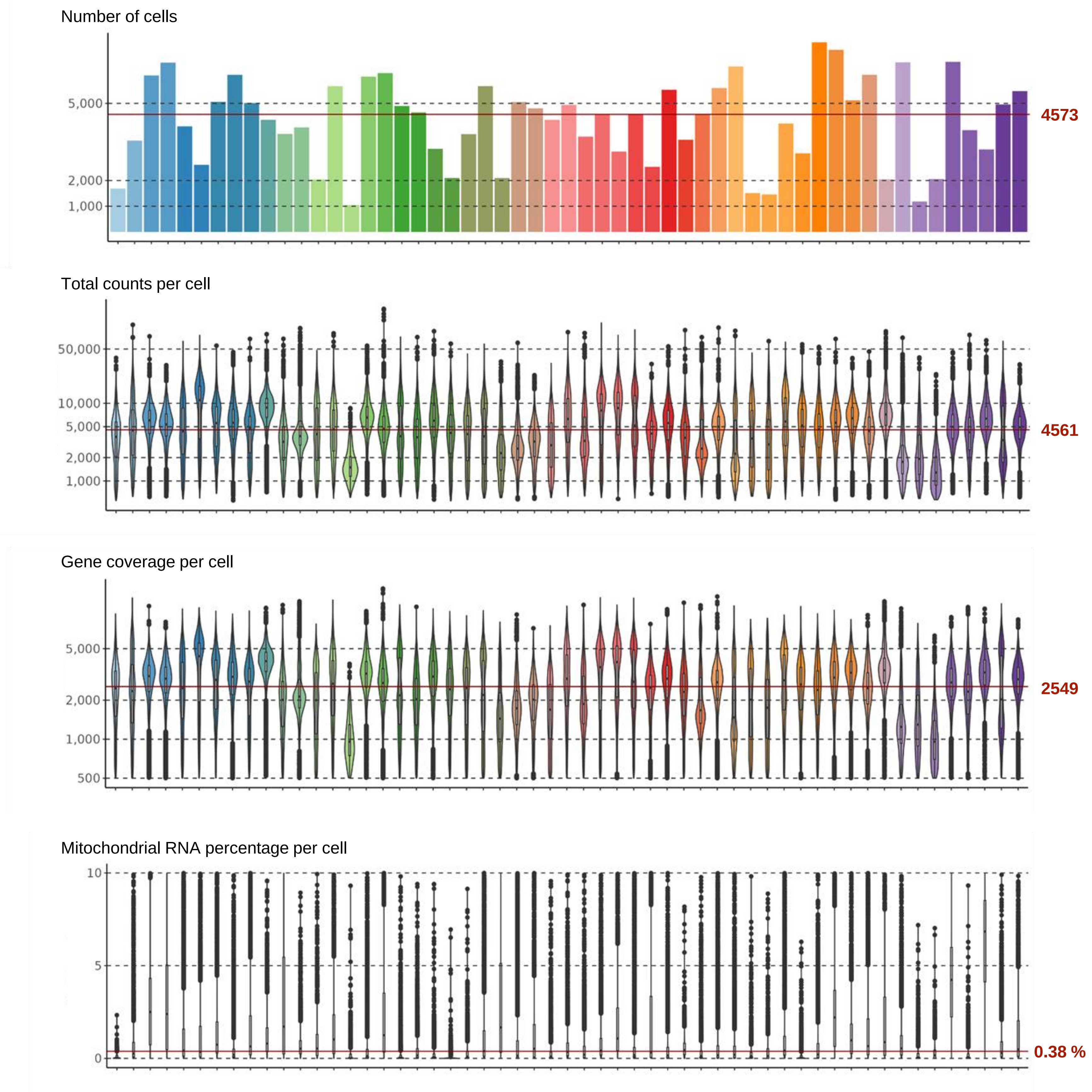


**Supplementary Figure S1.** Description of quality control metrics per sample (from top to bottom: number of cells in sample, total counts/cell in sample, gene coverage/cell in sample, mitochondrial RNA percentage/cell in sample). Red full lines highlight median values across the cohort for each metric.

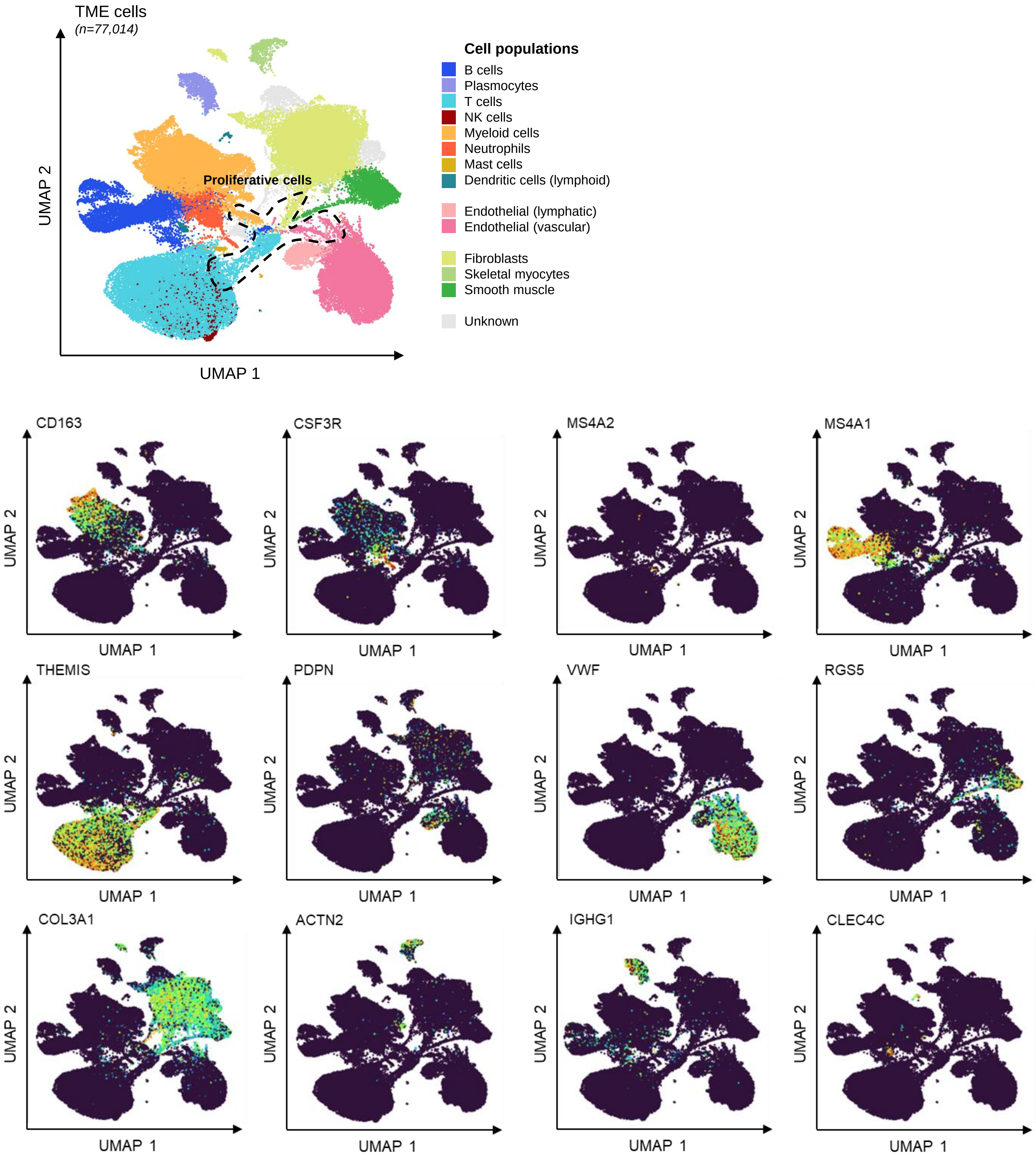


**Supplementary Figure S2.** Annotations of subpopulations in TME cells (top) and expression of correpsonding markers (bottom).

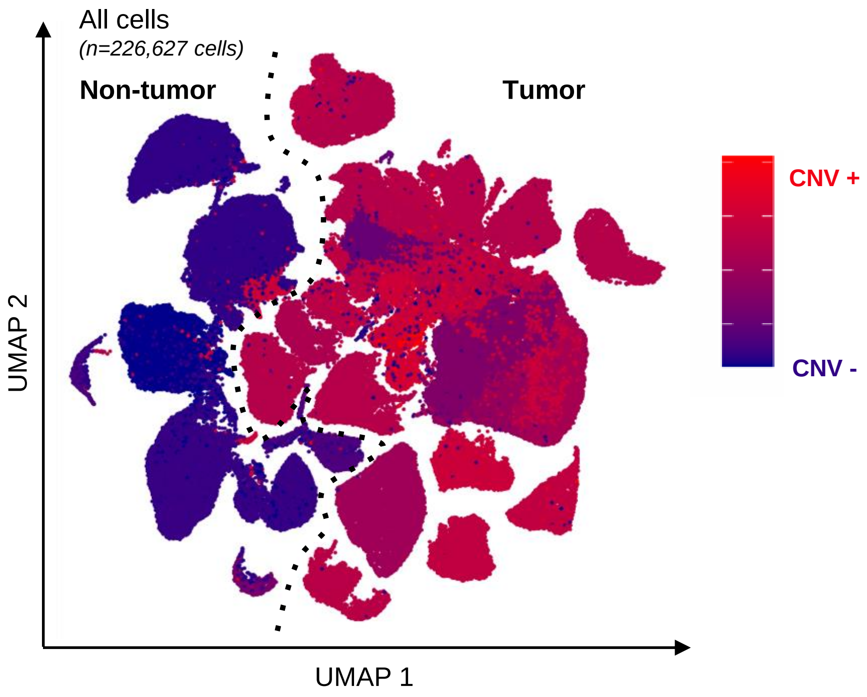


**Supplementary Figure S3.** UMAP of Copy Number Variation (CNV) scores generated by inferCNVpy projected in tumor cells and TME cells.

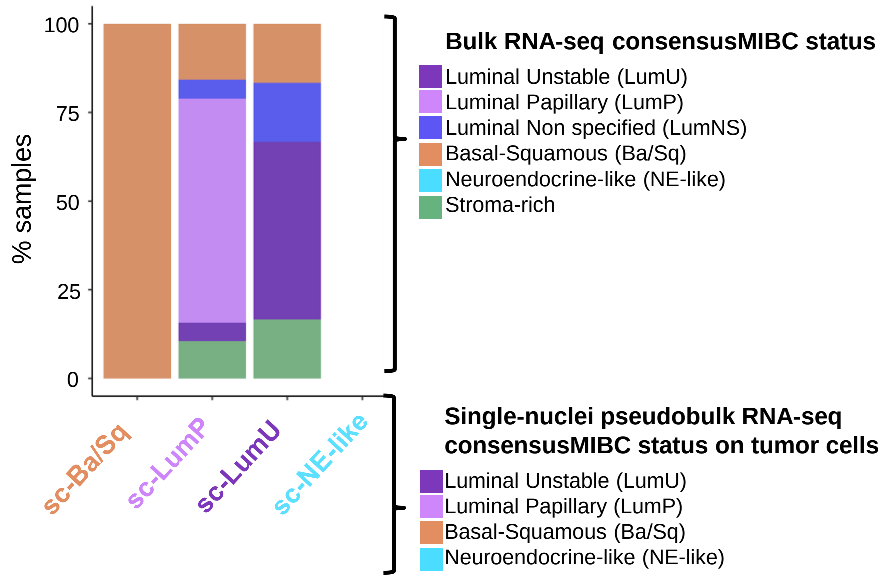


**Supplementary Figure S4.** Consensus classification (Kamoun et al. 2020) of samples according to pseudobulk single-nuclei sequencing and bulk transcriptome (N=30).

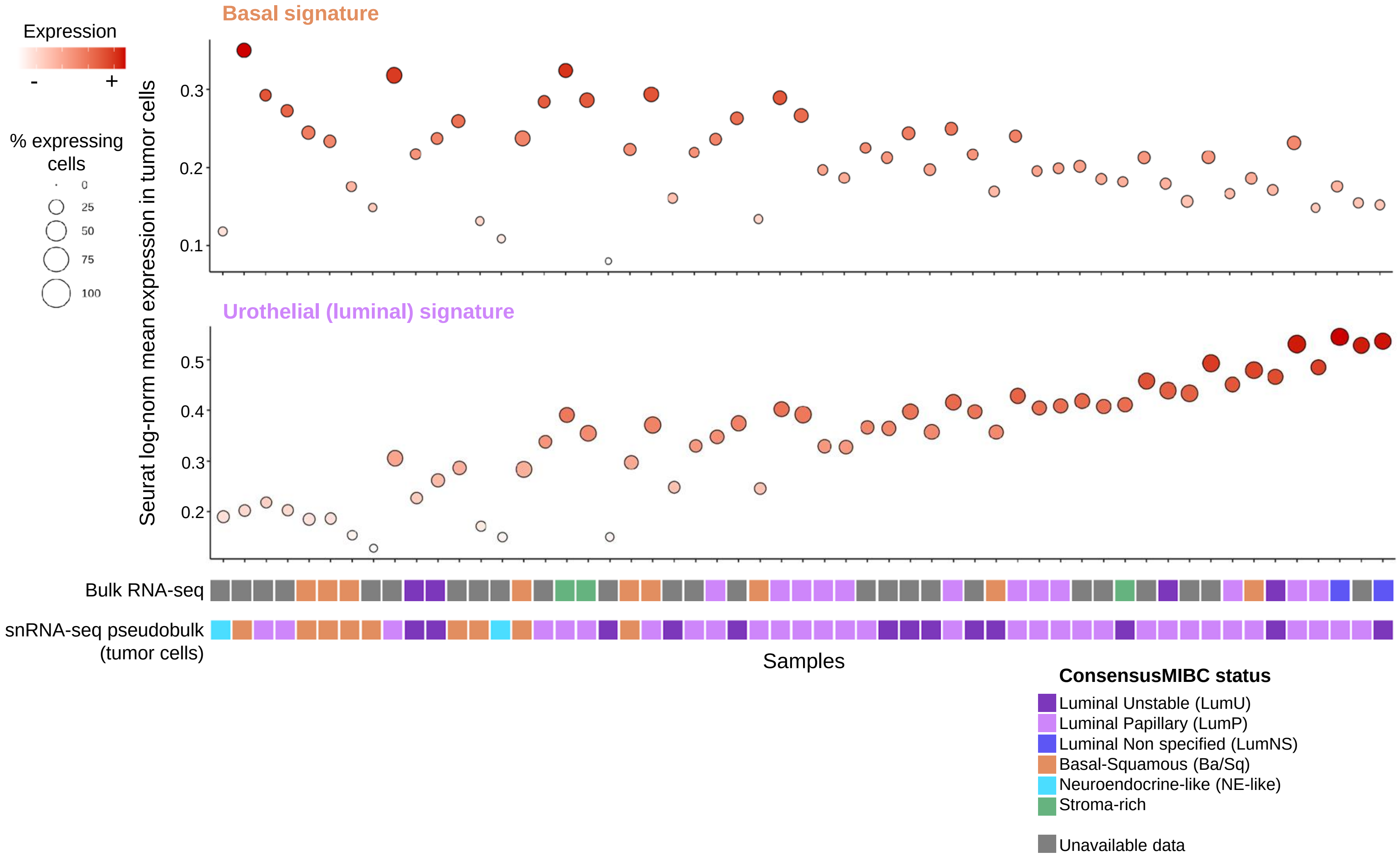


**Supplementary Figure S5.** Heterogeneity of single-nuclei tumor cell transcriptome by sample. Samples are ordered by mean expression difference between basal (n=143 genes) and urothelial (luminal, n=194 genes) differentiation signatures.

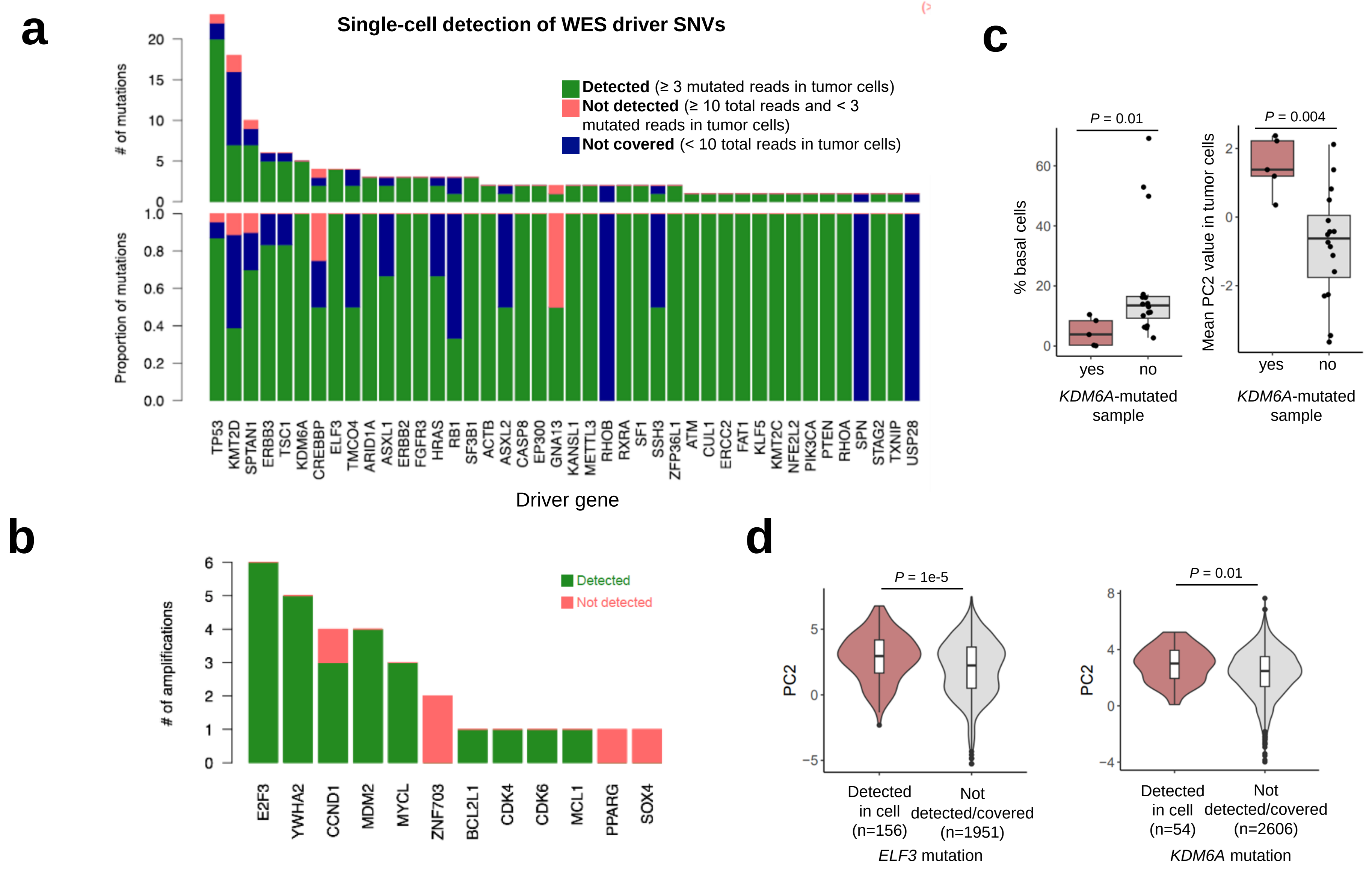


**Supplementary Figure S6.** Single-nuclei detection of driver alterations. **a.** Single-cell detection of driver mutations identified by whole exome sequencing. **b.** Single-cell detection of high-level amplifications identified by whole exome sequencing. **c.** *KDM6A*-mutated samples display significantly lower basal cell content and higher PC2 values. **d.** Cells in which *ELF3* and *KDM6A* mutations were detected display significantly higher PC2 values.

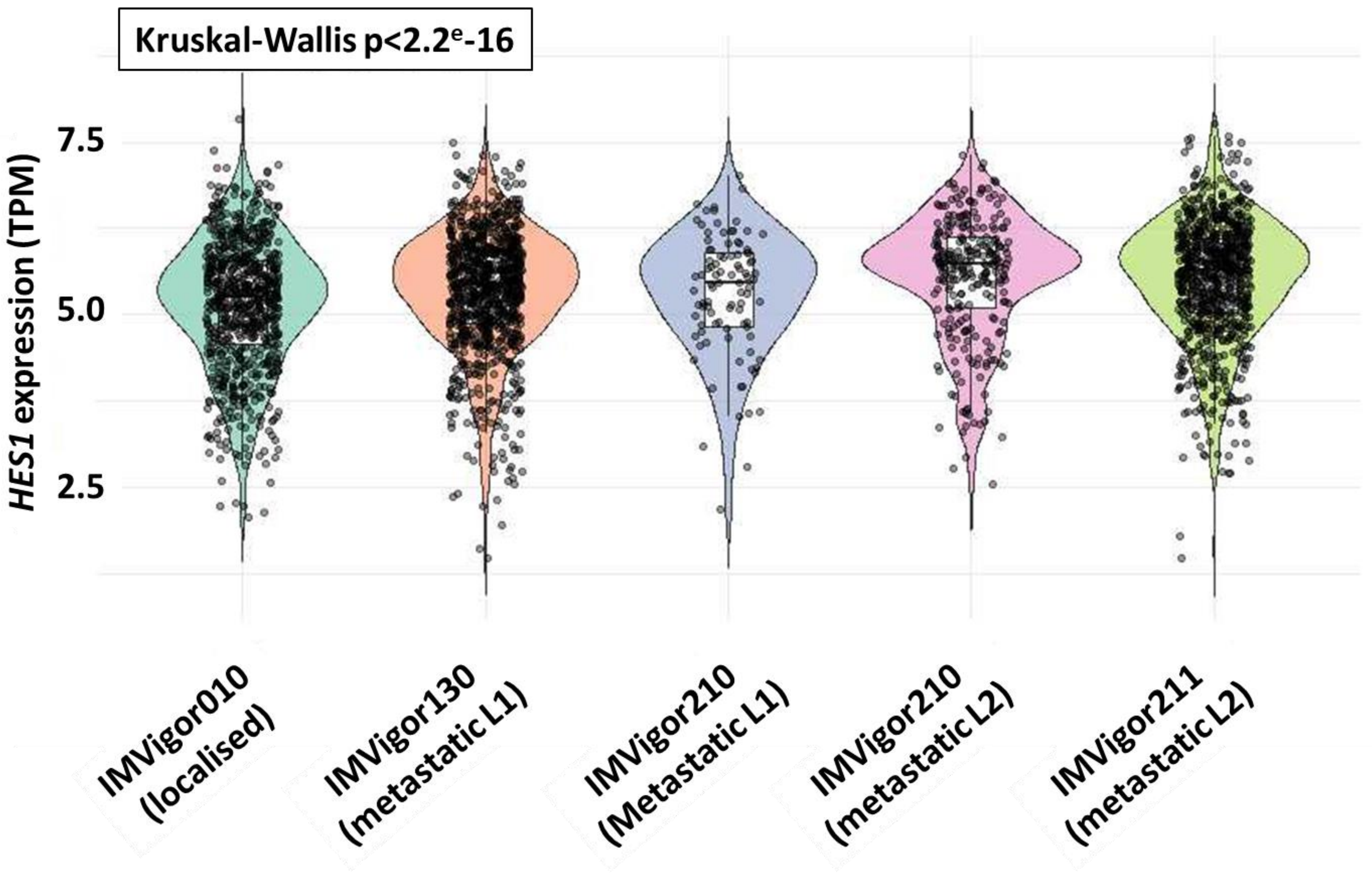


**Supplementary Figure S7.** Expression of HES1 across pivotal trials of atezolizumab in urothelial cancers across stages and lines of therapy.

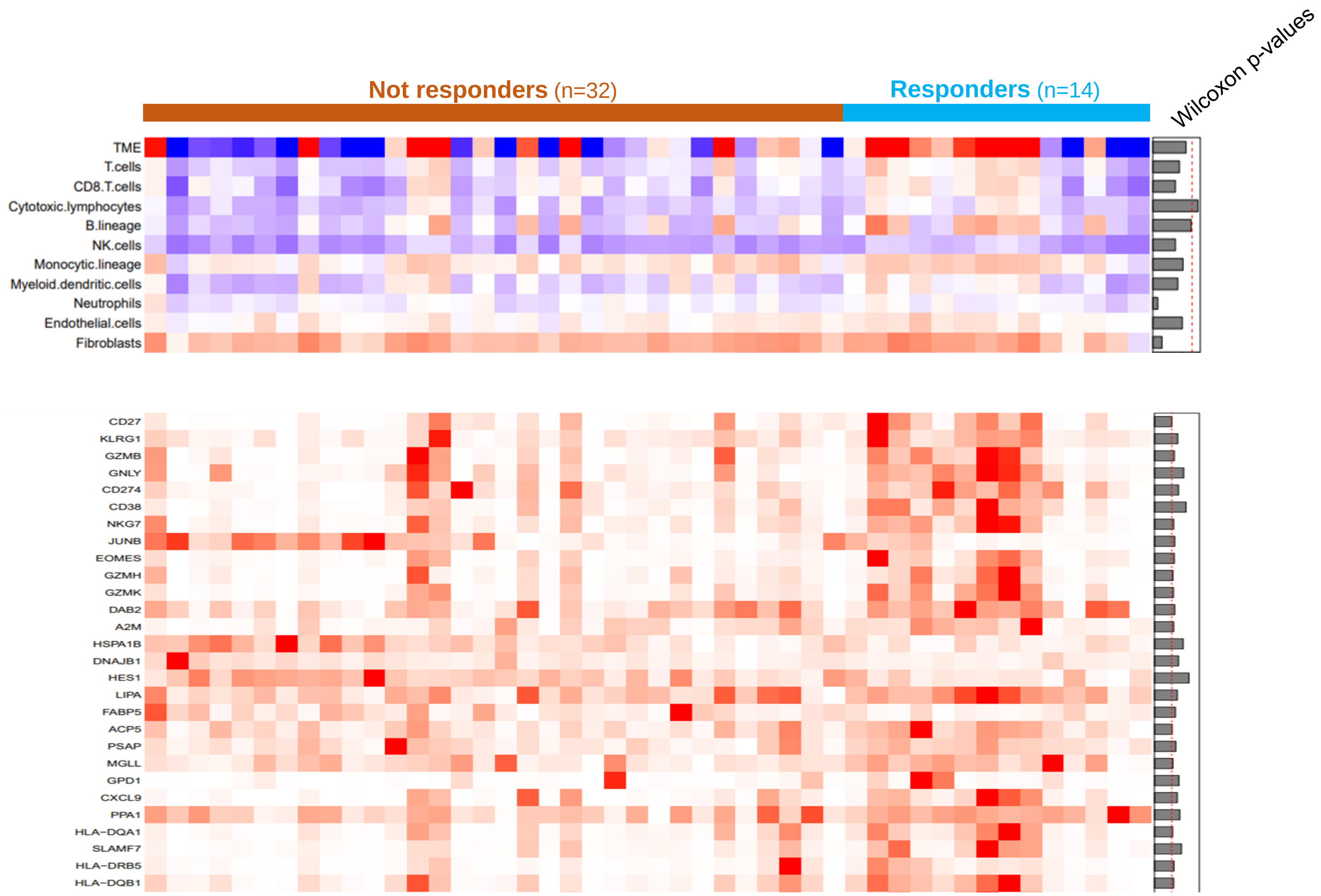


**Supplementary Figure S8.** (Top) Bulk RNA sequencing of the MATCH-R baseline cohort displaying inferred MCP-Counter scored immune cell types and (bottom) expression of immune-related genes according to response to therapy
